# Improved genetically encoded near-infrared fluorescent calcium ion indicators for *in vivo* imaging

**DOI:** 10.1101/2020.04.08.032433

**Authors:** Yong Qian, Danielle M. Orozco Cosio, Kiryl D. Piatkevich, Sarah Aufmkolk, Wan-Chi Su, Orhan T. Celiker, Anne Schohl, Mitchell H. Murdock, Abhi Aggarwal, Yu-Fen Chang, Paul W. Wiseman, Edward S. Ruthazer, Edward S. Boyden, Robert E. Campbell

**Author notes:** Correspondence and requests for materials should be addressed to R.E.C.

## Abstract

Near-infrared (NIR) genetically-encoded calcium ion (Ca^2+^) indicators (GECIs) can provide advantages over visible wavelength fluorescent GECIs in terms of reduced phototoxicity, minimal spectral cross-talk with visible-light excitable optogenetic tools and fluorescent probes, and decreased scattering and absorption in mammalian tissues. Our previously reported NIR GECI, NIR-GECO1, has these advantages but also has several disadvantages including lower brightness and limited fluorescence response compared to state-of-the-art visible wavelength GECIs, when used for imaging of neuronal activity. Here, we report two improved NIR GECI variants, designated NIR-GECO2 and NIR-GECO2G, derived from NIR-GECO1. We characterized the performance of the new NIR GECIs in cultured cells, acute mouse brain slices, and *C. elegans* and *Xenopus laevis in vivo*. Our results demonstrate that NIR-GECO2 and NIR-GECO2G provide substantial improvements over NIR-GECO1 for imaging of neuronal Ca^2+^ dynamics.

## Introduction

Fluorescence imaging of intracellular calcium ion (Ca^2+^) transients using genetically-encoded Ca^2+^ indicators (GECIs) is a powerful and effective technique to monitor *in vivo* neuron activity in model organisms^1–4^. Over a time frame spanning two decades^5,6^, tremendous effort has been invested in the development of visible wavelength GECIs based on green and red fluorescent proteins (GFP and RFP, respectively). These efforts have produced a series of high-performance GECIs that are highly optimized in terms of brightness, kinetics, Ca^2+^ affinities, cooperativity, and resting (baseline) fluorescence^2–4,7^. In contrast, efforts to develop GECIs with near-infrared (NIR) excitation and emission (>650 nm) are at a relatively nascent state^8,9^.

We recently described the conversion of a NIR FP (mIFP, a biliverdin (BV)-binding NIR FP engineered from the PAS and GAF domains of *Bradyrhizobium* bacteriophytochrome)^10^ into a GECI designated NIR-GECO1. NIR-GECO1 was engineered by genetic insertion of the Ca^2+^-responsive domain calmodulin (CaM)-RS20 into the protein loop close to the BV-binding site of mIFP^8^. NIR-GECO1 provides a robust inverted fluorescence response (that is, a fluorescence decrease upon Ca^2+^ increase) in response to Ca^2+^ concentration changes in cultured cells, primary neurons, and acute brain slices. Due to its NIR fluorescence, it has inherent advantages relative to visible wavelength GFP- and RFP-based GECIs including reduced phototoxicity, minimal spectral cross-talk with visible-light excitable optogenetic tools and fluorescent probes, and decreased scattering and absorption of excitation and emission light in mammalian tissues. However, as a first-generation GECI, NIR-GECO1 is suboptimal by several metrics including relatively low brightness and limited fluorescence response (that is, ΔF/F_0_ for a given change in Ca^2+^ concentration), which limit its utility for *in vivo* imaging of neuronal activity.

## Results

### Development of new NIR-GECO variants

Based on ample precedent from the development of the GFP-based GECI GCaMP series^3,6^ and the RFP-based R-GECO series^2,11^, we reasoned that NIR-GECO1 was likely to be amenable to further improvement by protein engineering and directed molecular evolution. Here we report that such an effort has led to the development of two second-generation variants, designated NIR-GECO2 and NIR-GECO2G, which enable fluorescence imaging of neuronal activity-associated changes in intracellular Ca^2+^ concentration with substantially greater sensitivity than their first-generation progenitor.

Starting from the template of the gene encoding NIR-GECO1, three rounds of directed evolution were performed as described previously^8^. Briefly, the gene encoding NIR-GECO1 was randomly mutated by error-prone PCR and the resulting gene library was used to transform *E. coli*. Petri dishes of colonies were screened using a fluorescence macro-imaging system, brightly fluorescent clones were picked and, following overnight culture, the NIR-GECO1 protein variants were extracted and tested for Ca^2+^ responsiveness *in vitro*. The genes encoding the best variants, as determined by the *in vitro* test, were assessed for brightness and Ca^2+^ responsiveness when expressed in HeLa cells. The gene that provided the best balance of brightness and function in HeLa cells was used as the template for a subsequent round of combined bacterial and HeLa cell screening. Three rounds of screening led to the identification of two promising variants: NIR-GECO2 (equivalent to NIR-GECO1 with T234I, S250T, E259G, Q402E, F463Y, and T478A, numbered as in **Supplementary Fig. 1**) and NIR-GECO2G (equivalent to NIR-GECO2 with T250S and S347G; **Fig. 1a**). In terms of fluorescence spectral profile, peak maxima, extinction coefficient, quantum yield, and pK_a_, NIR-GECO2 and NIR-GECO2G are essentially identical to NIR-GECO1 (**Supplementary Table 1**). One of the most pronounced changes in the biophysical properties is that the Ca^2+^ affinities of NIR-GECO2 and NIR-GECO2G are higher than that of NIR-GECO1 with *K*_d_ values of 331 nM and 480 nM, respectively (*K*_d_ of NIR-GECO1 is 885 nM) (**Supplementary Fig. 2a**). A parallel effort to construct a second-generation NIR-GECO1 by replacing the mIFP portion with the brighter and homologous miRFP^12^ resulted in the functional indicator prototype. However further optimization was abandoned due to the apparent toxicity of the miRFP-based construct when expressed in *E. coli* (**Supplementary Figs. 3,4**).

**Figure 1.**
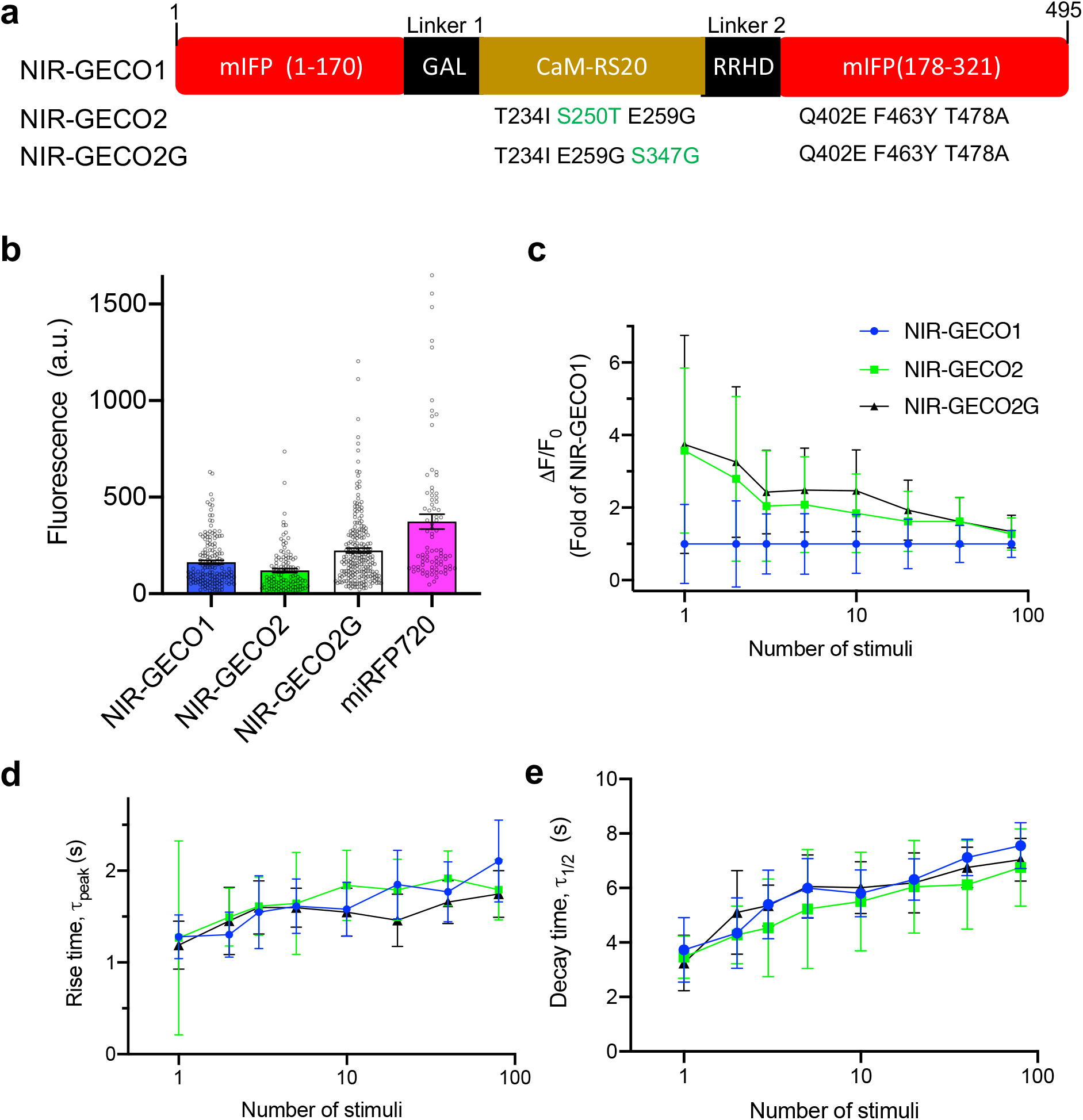
NIR-GECO evolution and characterization in dissociated neurons. (**a**) Mutations of NIR-GECO2 and NIR-GECO2G relative to NIRGECO1. The different mutations between NIR-GECO2 and NIR-GECO2G are highlighted in green. (**b**) Relative fluorescence intensity (mean ± s.e.m) of NIR-GECO1, NIR-GECO2, NIR-GECO2G and miRFP720 in neurons (n = 160, 120, 219, 84 neurons, respectively, from 2 cultures). Fluorescence was normalized by co-expression of GFP via self-cleavable 2A peptide. (**c-e**) Comparison of NIR-GECO variants, as a function of stimulus strength (the same color code is used in panel **c**-**e**). (**c**) ΔF/F_0_; (**d**) rise time; (**e**) half decay time. Values are shown as mean ± s.d. (n = 10 wells from 3 cultures).

### Characterization of new NIR-GECO variants

To compare the intracellular baseline brightness (that is, fluorescence in a resting state neuron) of NIR-GECO2 and NIR-GECO2G with the previously reported NIR-GECO1 and the NIR FP miRFP720 (Ref.13), we expressed each construct in cultured neurons and quantified the overall cellular brightness 5 days after transfection. To correct for cell-to-cell variations in protein expression, we co-expressed GFP stoichiometrically via a self-cleaving 2A peptide^14^ to serve as an internal reference for expression level. Under these conditions, NIR-GECO2 is approximately 25% dimmer than NIR-GECO1 while NIR-GECO2G is 50% brighter than NIR-GECO1 (**Fig. 1b**). Photobleaching experiments for the neuronally-expressed constructs revealed that all the NIR-GECO variants possess similar photostability and photobleach ~4× faster than miRFP720 (**Supplementary Fig. 2b**).

To assess the sensitivity (i.e., -ΔF/F_0_) of the new NIR-GECO variants, electric field stimulation^15^ was performed to cultured neurons expressing NIR-GECO2, NIR-GECO2G and NIR-GECO1. For a single field stimulation-evoked action potential (AP), NIR-GECO2 and NIR-GECO2G responded with similar -ΔF/F_0_ values of 16% and 17%, respectively. These values are 3.6- and 3.7-fold higher, respectively, than that of NIR-GECO1 (4.5%). For small numbers of APs (2 to 10), the responses of NIR-GECO2G were 2.5-to 3.3-fold larger than those of NIR-GECO1 and the responses of NIR-GECO2 were 1.8- to 2.8-fold larger than those of NIR-GECO1. At higher numbers of APs (20 to 80), the improvements of the new variants became less pronounced as the ΔF/F_0_ values of the three variants converged (**Fig. 1c**). The on (rise time, τ_peak_; **Fig. 1d**) and off (decay time, τ_1/2_; **Fig. 1e**) kinetics of NIR-GECO2 and NIR-GECO2G, in response to field stimulation-evoked APs stimuli, remained similar to that of NIR-GECO1.

To investigate if NIR-GECO2 and NIR-GECO2G provide advantages over NIR-GECO1 for combined use with an optogenetic actuator, we co-transfected HeLa cells with the genes encoding Opto-CRAC and each of the three NIR-GECO variants. Opto-CRAC is an optogenetic tool that can be used to induce Ca^2+^ influx into non-excitatory cells when illuminated with blue light^16^. Transfected HeLa cells were illuminated with 470 nm light at a power of 1.9 mW/mm^2^ while the NIR fluorescence intensity of NIR-GECO variants was continuously recorded. Following 100 ms of blue light stimulation, the average -ΔF/F_0_ for NIR-GECO2G, NIR-GECO2, and NIR-GECO1 was 34.5%, 22.8%, and 12.1%, respectively (**Fig. 2a, d**). With 500 ms of illumination, the -ΔF/F_0_ values increased to 48.2%, 42.1% and 30.7%, respectively. At 1 s of illumination time, the -ΔF/F_0_ was 40.7%, 38.0%, and 33.3%, respectively (**Fig. 2a, b, c**). When expressed in acute brain slices, both NIR-GECO2 and NIR-GECO2G robustly reported Ca^2+^ changes in neurons in response to either optogenetic (CoChR) or chemical (4-aminopyridine) stimulation (**Supplementary Fig. 5**). These results support the conclusion that both NIR-GECO2G and NIR-GECO2 are more sensitive than NIR-GECO1 for reporting Ca^2+^ transients at low Ca^2+^ concentrations, and NIR-GECO2G is the best of the three. To further explore the use of NIR-GECO2G, we expressed it in human induced pluripotent stem cell-derived cardiomyocytes (iPSC-CMs). In iPSC-CMs, NIR-GECO2G enabled robust imaging of spontaneous Ca^2+^ oscillations, caffeine induced Ca^2+^ influx, and channelrhodopsin-2 (ChR2)-dependent activation (**Supplementary Fig. 6**).

**Figure 2.**
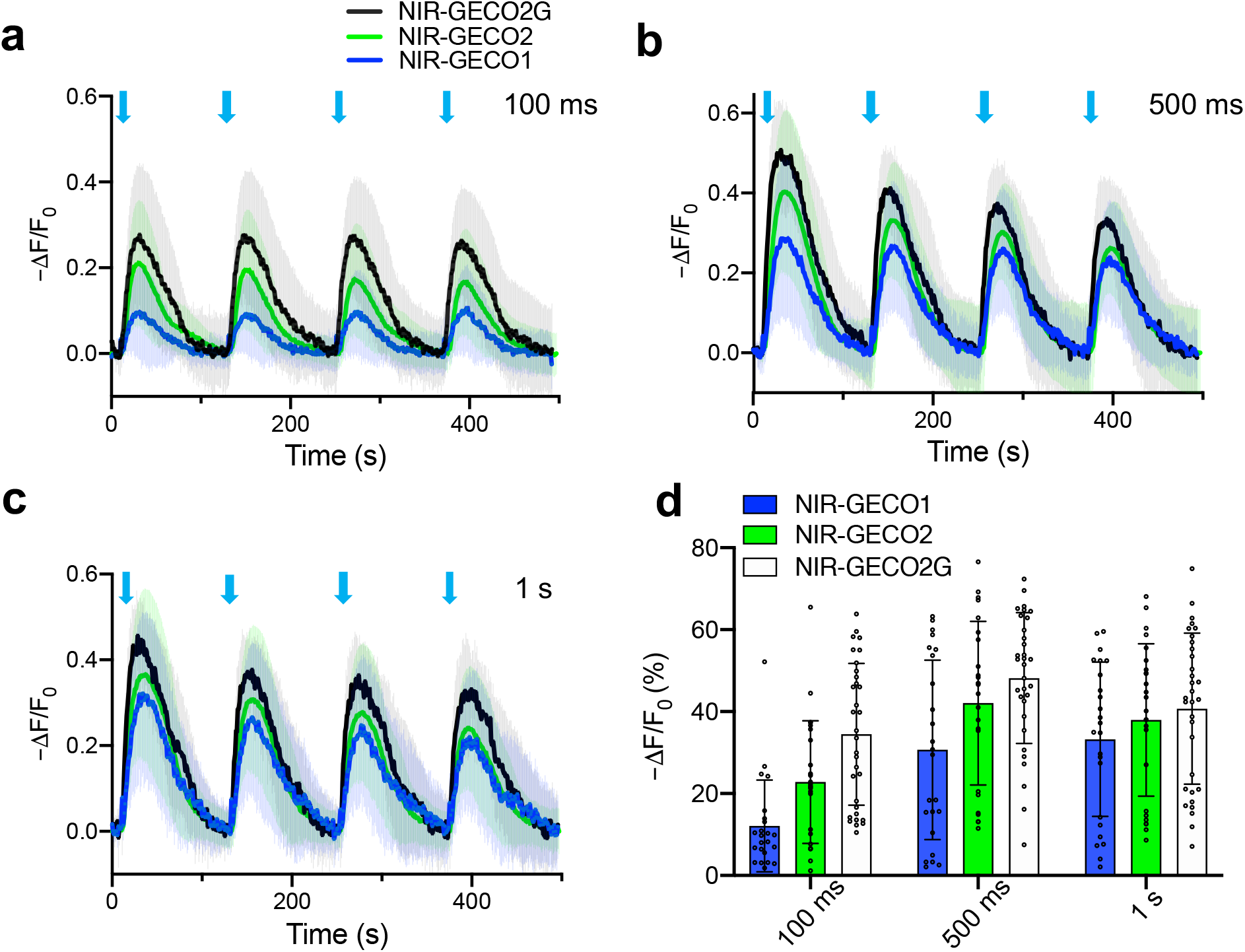
Performance of NIR-GECO2G, NIR-GECO2 and NIR-GECO1 in HeLa cells. (**a**-**c**) Fluorescence traces of NIR-GECO2G, NIR-GECO2 and NIR-GECO1 in response to 100 ms (**a**), 500 ms (**b**), and 1 s (**c**) blue light activation (470 nm at a power of 1.9 mW/mm^2^) in HeLa cells with co-expression of Opto-CRAC. Opto-CRAC is composed of the STIM1-CT and LOV2 domain. The fusion of STIM1-CT to the LOV2 domain allows photo-controllable exposure of the active site of SRIMI-CT, which is able to interact with ORAI1 and trigger Ca^2+^ entries across the plasma membrane^16^. Black, green and dark blue line represent averaged data for NIR-GECO2G (n = 32 cells), NIR-GECO2 (n = 26 cells) and NIR-GECO1(n = 23 cells), respectively. The same color code is used in panel **a-c**. Shaded areas represent the standard deviation. (**d**) Quantitative -ΔF/F_0_ (mean ± s.d.) for NIR-GECO2G, NIR-GECO2 and NIR-GECO1 in **a-c**.

### *In vivo* imaging of Ca^2+^ in *C. elegans* using NIR-GECO2

To determine if NIR-GECO2 was suitable for *in vivo* imaging of neuronal activity, we first sought to test it in *Caenorhabditis elegans*, a popular model organism in neuroscience. For this application, we chose to use NIR-GECO2 rather than NIR-GECO2G due to its higher Ca^2+^ affinity, however NIR-GECO2G could be also readily expressed in *C.elegans* in neurons producing sufficient near-infrared fluorescence (**Supplementary Figure 7d**). As the internal BV concentration of *C. elegans* is quite low due to its inability to synthesize heme *de novo* (its main source of heme is from the ingestion of *E. coli*)^17,18^, we decided to coexpress heme-oxygenase1 (HO1) to increase the conversion of heme into BV^19^. We created *C. elegans* lines expressing NLS-NIR-GECO2-T2A-HO1 (where NLS is a nuclear localization sequence) and NLS-jGCaMP7s under the pan-neuronal tag-168 promoter in an extrachromosomal array. The resulting transgenic worms exhibited bright nuclear localized fluorescence from both NIR-GECO2 and jGCaMP7s. One notable advantage of the NIR-GECO series relative to the GCaMP series of indicators was the lower auto-fluorescence in the intestinal area of worms in the NIR fluorescence channel, as compared to the green fluorescence channel (**Fig. 3a, Supplementary Fig. 7a,d**). Microfluidic chips^20^ were used to deliver a high-osmotic-strength stimulus (200 mM NaCl) to individual worms, and the fluorescence was imaged simultaneously in the NIR and green fluorescence channels. Following exposure to a high concentration of NaCl, we detected synchronous but opposing fluorescent changes for jGCaMP7s (fluorescence increases) and NIR-GECO2 (fluorescence decreases)(**Fig. 3b**). Quantitative analysis of 36 spikes from 3 neurons showed that the -ΔF/F_0_ of NIR-GECO2 was about half of the ΔF/F_0_ of jGCaMP7s following NaCl stimulation (ΔF/F_0_ = 0.39 ± 0.19 for jGCaMP7s; -ΔF/F_0_ = 0.19 ± 0.07 for NIR-GECO2; **Fig. 2c**).

**Figure 3.**
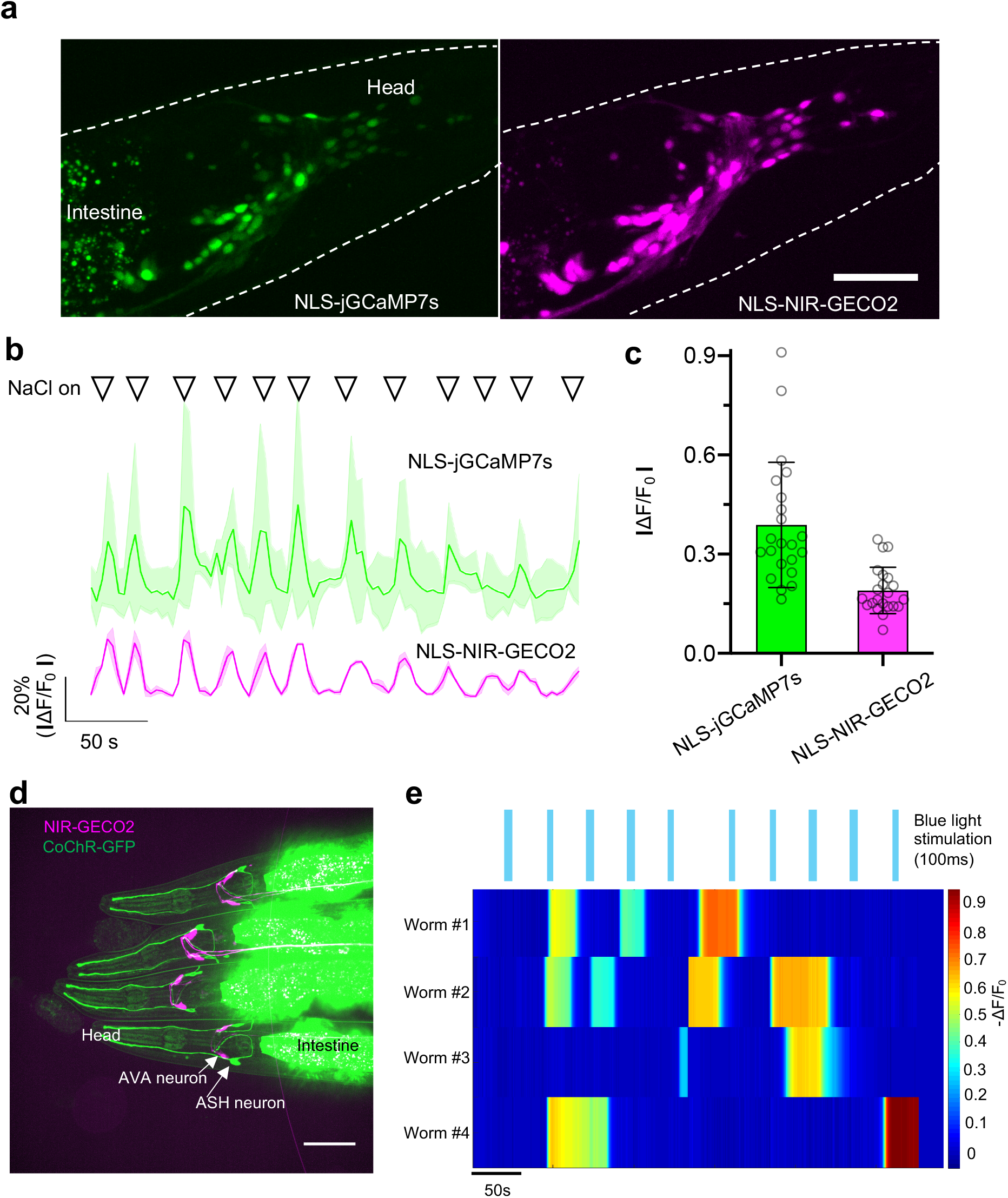
Imaging of NIR-GECO2 in response to microfluidic and optogenetic stimulation in *C. elegans in vivo*. (**a**) Left, fluorescent image of neurons expressing NLS-jGCaMP7s (λ_ex_ = 488 nm laser light, λ_em_ = 527/50 nm). Right, fluorescent image of neurons expressing NIR-GECO2-T2A-HO1 (λ_ex_ = 640 nm laser light, λ_em_ = 685/40 nm). Representative of more than 3 worms, both under tag-168 promoter. Scale bar, 50 μm. (**b**) Fluorescence traces of NLS-jGCaMP7s (top) and NLS-NIR-GECO2 (bottom) in response to the stimulation of microfluidic containing 200mM of NaCl. Solid lines represent averaged data from 3 neurons. Shaded areas are shown as standard deviation. Triangles on the top of the traces indicate the time points of stimulation (20 s for each stimulation). (**c**) Quantitative fluorescence changes of NLS-jGCaMP7s and NLS-NIR-GECO2 in **b**(n = 36 spikes from 3 neurons). (**d**) Fluorescence image of the 4 *C. elegans* expressing NIR-GECO2-T2A-HO1 in AVA neurons (under flp-18 promoter) and CoChR-GFP in ASH neurons (under sra-6 promoter). The merged image is shown. Imaging condition: NIR-GECO2, λ_ex_ = 637 nm laser light, λ_em_ = 664LP; GFP, λ_ex_ = 488 nm laser light, λ_ex_=527/50 nm. (**e**) Individual traces of NIR-GECO2 fluorescence in an AVA neuron under blue light illumination (20 mW/mm^2^, λ_ex_ = 488 nm laser light, 100 ms; blue bars).

We next attempted all-optical stimulation and imaging of neuron activity in *C. elegans* using the blue-light sensitive channelrhodopsin CoChR^21^ and NIR-GECO2. We previously demonstrated that excitation wavelengths used to image NIR-GECO1 do not activate CoChR^8^. NIR-GECO2 (with co-expression of HO1) was expressed in AVA interneurons (involved in backward locomotion) under the flp-18 promoter, and CoChR-GFP was expressed in upstream ASH neurons under control of the sra-6 promoter. Imaging of transgenic worms with confocal microscopy revealed two AVA neurons with expression of NIR-GECO2 and two ASH neurons with expression of CoChR (**Fig. 3d**). Blue-light stimulation of CoChR in ASH neurons caused long-lasting (tens of seconds to a few minutes) fluorescent decreases in NIR-GECO2 fluorescence (-ΔF/F_0_ of 30% to 90%) in the downstream AVA interneurons (**Fig. 3e**). Collectively, this data indicates that the combination of NIR-GECO2 and CoChR provides a robust all-optical method to interrogate hierarchical circuits in *C. elegans*.

### *In vivo* imaging of Ca^2+^ in *Xenopus laevis* using NIR-GECO2G

To further evaluate the utility of NIR-GECO2G for *in vivo* imaging of neuronal activity in a vertebrate brain, we transiently expressed the genes encoding NIR-GECO2G (without co-expression of HO1) and GCaMP6s^1^ in *Xenopus laevis* tadpoles by mRNA injection into early embryos. Light-sheet microscopy imaging of the olfactory bulb of live tadpoles revealed that individual neurons exhibited strong near-infrared fluorescent signals due to NIR-GECO2G expression (**Fig. 4a, b, Supplementary Video 1**). Fluorescence from both NIR-GECO2G and GCaMP6s was observed to oscillate in a synchronous but opposing manner in response to spontaneous neuronal activity (**Fig. 4c**). These results demonstrate that NIR-GECO2G can be used to report dynamic Ca^2+^ changes *in vivo* in *Xenopus laevis* in the absence of added BV or HO1 co-expression.

**Figure 4.**
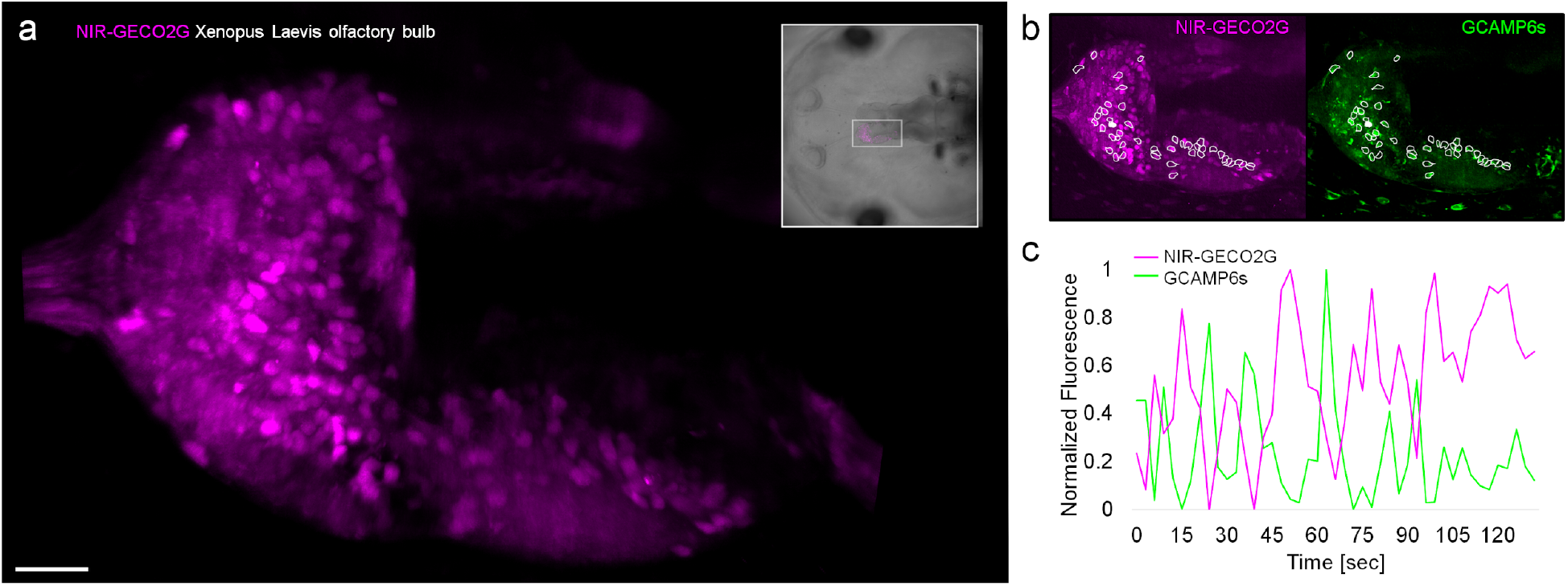
Spontaneous Ca^2+^ response imaging with NIR-GECO2G in the olfactory bulb of *Xenopus laevis*. (**a**) Light-sheet acquisition of spontaneous Ca^2+^ response in the olfactory bulb in an intact animal. The image shows an intensity projection over 200 frames and a volume of 45 μm. The orientation and position of the field of view is marked in the brightfield image of the animal (upper right frame). Scalebar 40 μm. (**b**) White frames indicate the cells that showed Ca^2+^ response with the NIR-GECO2G (magenta) and GCAMP6s (green) indicator. (**c**) A representative fluorescence response trace of spontaneous activity of one cell in the olfactory bulb (solid white ROI in **b**) is plotted over time. The intensity responses of NIR-GECO2G and GCAMP6s are antagonal to each other. Z-series volumes were acquired once every 3.0 s.

## Discussion

In summary, we have developed two improved NIR fluorescent Ca^2+^ indicators designated NIR-GECO2 and NIR-GECO2G. Of the two, NIR-GECO2 has higher response amplitudes but dimmer fluorescence compared to NIR-GECO1, based on characterization in neurons and HeLa cells. In contrast, NIR-GECO2G is improved relative to NIR-GECO1 in terms of both overall cellular brightness (~50% brighter than NIR-GECO1) and sensitivity (up to a ~3.7 fold improvement in -ΔF/F_0_ relative to NIR-GECO1 for single AP). As we have demonstrated in this work, these improvements make the new variants particularly useful for imaging Ca^2+^ dynamics in small model organisms. Specifically, NIR-GECO2 offers comparable sensitivity to jGCaMP7s in *C. elegans* and NIR-GECO2G enables robust imaging of Ca^2+^ dynamics in the olfactory bulb of *Xenopus laevis* tadpoles. However, even with these improvements, NIR GECIs still face challenges like relatively low brightness, slower kinetics, and faster photobleaching compared to the state-of-art green and red fluorescent GECIs. Overcoming these challenges will undoubtedly require further directed molecular evolution and optimization of the NIR-GECO series, or the possible development of alternative NIR GECI designs based on brighter and more photostable NIR FP scaffolds.

## Methods and Materials

### Mutagenesis and molecular cloning

Synthetic DNA oligonucleotides used for cloning and library construction were purchased from Integrated DNA Technologies. Random mutagenesis of NIR-GECO variants was performed using *Taq* DNA polymerase (New England BioLabs) with conditions that resulted in a mutation frequency of 1-2 mutations per 1,000 base pairs. Gene fragments for NIR-GECO libraries were then inserted between restriction sites *XhoI* and *HindIII* of pcDuex2 for expression. The DNA sequences encoding miRFP_1 to 172_, CaM-RS20 (from NIR-GECO1), and miRFP_179 to 311_ were amplified by PCR amplification separately and then used as DNA templates for the assembly of miRFP_1 to 172_ - CaM-RS20-miRFP_179 to 311_ by overlap extension PCR. The resulting DNA sequence was then digested and ligated into the pcDNA3.1(-) vector for mammalian expression and into a pBAD-MycHisC (Invitrogen) vector for bacterial expression. Q5 high-fidelity DNA polymerase (New England BioLabs) was used for routine PCR amplifications and overlap extension PCR. PCR products and products of restriction digests were routinely purified using preparative agarose gel electrophoresis followed by DNA isolation with the GeneJET gel extraction kit (Thermo Fisher Scientific). Restriction endonucleases were purchased from Thermo Fisher Scientific. Ligations were performed using T4 ligase in Rapid Ligation Buffer (Thermo Fisher Scientific).

Genes or gene libraries in expression plasmids were electroporated into *E. coli* strain DH10B (ThermoFisher Scientific). The transformed cells were then plated on 10 cm Lysogeny broth (LB)-agar Petri dishes supplemented with 400 μg/mL ampicillin (Sigma) and 0.0004% (wt/vol) L-arabinose (Alfa Aesar) at 37 °C overnight. For library screening, bright bacterial colonies expressing NIR-GECO variants were picked and cultured. Proteins were extracted using B-PER (ThermoFisher Scientific) from overnight cultures of bacteria growing in LB media supplemented with 100 μg/mL ampicillin and 0.0016% L-arabinose and then tested for fluorescence and Ca^2+^ response in 384-well plates. Variants with the highest brightness and Ca^2+^ response were selected and the corresponding plasmids were purified. HeLa cells were transfected with the selected plasmids and live-cell fluorescence imaging was used to re-evaluate both brightness and Ca^2+^ response. Small-scale isolation of plasmid DNA was done with GeneJET miniprep kit (Thermo Fisher Scientific).

For the construction of Opto-CRAC-EYFP, a synthetic double-stranded DNA fragment consisting of fused EYFP, LOV2 and STIM1-CT fragments (residues 336-486)^16^, flanked with *NotI* and *XhoI* restriction sites, was cloned into the pcDNA3.1(-) vector.

### Protein purification and *in vitro* characterization

The genes for the miRFP-based Ca^2+^ indicators NIR-GECO1, NIR-GECO2 and NIR-GECO2G, with a poly-histidine tag on the C-terminus, were expressed from a pBAD-MycHisC (Invitrogen) vector containing the gene of cyanobacteria *Synechocystis* HO1 as previously described^22,23^. Bacteria were lysed with a cell disruptor (Constant Systems Ltd) and then centrifuged at 15,000*g* for 30 min, and proteins were purified by Ni-NTA affinity chromatography (Agarose Bead Technologies). The buffer wasexchanged to 10 mM MOPS, 100 mM KCl (pH 7.2) with centrifugal concentrators (GE Healthcare Life Sciences). The spectra of miRFP-based Ca^2+^ indicator prototype, with and without Ca^2+^, were measured in a 384-well plate. Briefly, purified proteins were loaded into 384-well plates and then supplied with either 10 mM EGTA or 5 mM CaCl_2_ before measuring emission spectra. The extinction coefficients (EC), quantum yield (QY) and p*K*_a_ for NIR-GECO variants were determined as previously described^8^. Ca^2+^ titrations of NIR-GECO variants were performed with EGTA-buffered Ca^2+^ solutions. We prepared buffers by mixing a CaEGTA buffer (30 mM MOPS, 100 mM KCl, 10 mM EGTA, 10 mM CaCl_2_) and an EGTA buffer (30 mM MOPS, 100 mM KCl, 10 mM EGTA) to give free Ca^2+^ concentrations ranging from 0 nM to 39 μM at 25 °C. Fluorescence intensities were plotted against Ca^2+^ concentrations and fitted by a sigmoidal binding function to determine the Hill coefficient and *K*_d_. To determine k_off_ for NIR-GECO variants, a SX20 stopped-flow spectrometer (Applied Photophysics) was used. Proteins samples with 10 μM CaCl_2_ (30 mM MOPS, 100 mM KCl, pH 7.2) were rapidly mixed with 10 mM EGTA (30 mM MOPS, 100 mM KCl, pH 7.2) at room temperature, fluorescence growth curve was measured and fitted by a single exponential equation.

### Imaging of miRFP-based Ca^2+^ indicator prototype in HeLa cells

HeLa cells (40-60% confluent) on 24-well glass bottom plate (Cellvis) were transfected with 0.5 μg of plasmid DNA and 2 μl TurboFect (Thermo Fisher Scientific). Following 2 hours incubation, the media was changed to DEME (Gibco Fisher Scientific) with 10% fetal bovine serum (FBS) (Sigma), 2 mM GlutaMax (Thermo Fisher Scientific) and 1% penicillin-streptomycin (Gibco) and the cells were incubated for 48 hours at 37 °C in a CO_2_ incubator before imaging. Prior to imaging, culture medium was changed to HBSS. Wide-field imaging was performed on a Nikon Eclipse Ti microscope that was equipped with a 75 W Nikon xenon lamp, a 16-bit 512SC QuantEM EMCCD (Photometrics), and a 60× objective and was driven by a NIS-Elements AR 4.20 software package (Nikon). For time-lapse imaging, HeLa cells were treated with 4 mM EGTA (with 5 μM ionomycin) and then 10 mM CaCl_2_ (with 5 μM ionomycin). Images were taken every 5 seconds using a filter set with 650/60 nm excitation and 720/60 nm emission.

### Imaging of NIR-GECO1 and NIR-GECO2G in HeLa cells with Opto-CRAC

HeLa cells were co-transfected with pcDNA3.1-NIR-GECO1 or pcDNA3.1-NIR-GECO2 and pcDNA3.1-Opto-CRAC-EYFP using transfection reagent Lipofectamine 3000 (Invitrogen) following the manufacturer’s instructions. An inverted microscope (D1, Zeiss) equipped with a 63× objective lens (NA 1.4, Zeiss) and a multiwavelength LED light source (pE-4000, CoolLED) was used. Blue (470 nm) and red (635 nm) excitations were used to illuminate Opto-CRAC-EYFP and image NIR-GECO variants, respectively. The GFP filter set (BP 470-490, T495lpxr dichroic mirror, and HQ525/50 emission filter) and the NIR filter set (ET 650/45x, T685lpxr dichroic mirror, and ET720/60 emission filter) was used to confirm the expression of Opto-CRAC-EYFP and NIR-GECO variants. The filter set (T685lpxr dichroic mirror, and ET720/60 emission filter) was used to stimulate Opto-CRAC and to acquire fluorescence imaging of NIR-GECO1 and NIR-GECO2G. Optical stimulation was achieved with the 470 nm LED light at a power density of 1.9 mW/mm^2^. Fluorescence signals were recorded by a CMOS camera (ORCA-Flash4.0LT, Hamamatsu) and controlled by a software (HC Image).

### Imaging of NIR-GECO2G in Human iPSC-derived cardiomyocytes

Human iPSC-derived cardiomyocytes (human iPSC cardiomyocytes - male | ax2505) were purchased from Axol Bioscience. The 96 well glass-bottom plate was first coated with fibronectin and gelatin (0.5% and 0.1%, respectively) at 37 °C for at least 1 hour. The cells were then plated and cultured for three days in Axol’s Cardiomyocyte Maintenance Medium. IPSC-CMs were then transfected with pcDNA3.1-NIR-GECO2 with or without pcDNA3.1-hChR2-EYFP using Lipofectamine 3000 (Invitrogen) following the manufacturer’s instructions.The medium was switched to Tyrode’s buffer right before imaging. Imaging was performed with an inverted microscope (D1, Zeiss) equipped with a 63× objective lens (NA 1.4, Zeiss) and a multi-wavelength LED light source (pE-4000, CoolLED) using the same settings described above.

### Imaging of NIR-GECO1, NIR-GECO2, NIR-GECO2G and miRFP720 in cultured neurons

For dissociated hippocampal mouse neuron culture preparation, postnatal day 0 or 1 Swiss Webster mice (Taconic Biosciences, Albany, NY) were used as previously described^21^. Briefly, dissected hippocampal tissue was digested with 50 units of papain (Worthington Biochem) for 6-8 min at 37 °C, and the digestion was stopped by incubating with ovomucoid trypsin inhibitor (Worthington Biochem) for 4 min at 37 °C. Tissue was then gently dissociated with Pasteur pipettes, and dissociated neurons were plated at a density of 20,000–30,000 per glass coverslip coated with Matrigel (BD Biosciences). Neurons were seeded in 100 μL plating medium containing Minimum Essential Medium (MEM) (Thermo Fisher Scientific), glucose (33 mM, Sigma), transferrin (0.01%, Sigma), HEPES (10 mM, Sigma), Glutagro (2 mM, Corning), insulin (0.13%, Millipore), B27 supplement (2%, Gibco), and heat-inactivated FBS (7.5%, Corning). After cell adhesion, additional plating medium was added. AraC (0.002 mM, Sigma) was added when glia density was 50–70% of confluence. Neurons were grown at 37 °C and 5% CO_2_ in a humidified atmosphere.

To express each of NIR-GECO variants in primary hippocampal neurons and compare their brightness and photostability, the genes of NIR-GECO(1,2,2G)-T2A-GFP and were constructed using overlap-extension PCR followed by subcloning into pAAV-CAG vector (Addgene no. 108420) using BamHI and EcoRI sites. The gene for miRFP720-P2A-GFP was synthesized *de novo* by GenScript, based on the reported sequence^13^, and cloned into the pAAV-CAG vector. Cultured neurons were transfected with plasmids (1.5 μg of plasmid DNA per well) at 4 - 5 d in vitro (DIV) using a commercial calcium phosphate transfection kit (Thermo Fisher Scientific) as previously described^21,24^.

Wide-field fluorescence microscopy of cultured neurons was performed using an epifluorescence inverted microscope (Eclipse Ti-E, Nikon) equipped with an Orca-Flash4.0 V2 sCMOS camera (Hamamatsu) and a SPECTRA X light engine (Lumencor). The NIS-Elements Advanced Research (Nikon) was used for automated microscope and camera control. Cells were imaged with a 20× NA 0.75 air objective lens (Nikon) at room temperature for quantification of brightness or a 40× NA 1.15 for photobleaching experiments. Excitation: 631/28 nm; emission: 664LP.

### Field stimulation

Neurons expressing NIR-GECO variants (driven by CAG promoter) were imaged and stimulated in 24-well plates with 300 μL growth medium in each well at room temperature. Field stimuli (83 Hz, 50 V, 1 ms) were delivered in trains of 1, 2, 3, 5, 10, 20, 40, 80 via two platinum electrodes with a width of 6.5 mm to neurons. Neurons were imaged simultaneously while delivering trains of field stimuli with a 40× NA 1.15, a 631/28 nm LED (Spectra X light engine, Lumencor). Fluorescence was collected through 664LP using a sCMOS camera (Orca-Flash4.0, Hamamatsu) at 2 Hz.

### Animal care

All experimental manipulations performed at MIT were in accordance with protocols approved by the Massachusetts Institute of Technology Committee on Animal Care, following guidelines described in the US National Institutes of Health Guide for the Care and Use of Laboratory Animals. All procedures performed at McGill University were in accordance with the Canadian Council on Animal Care guidelines for the use of animals in research and approved by the Montreal Neurological Institute Animal Care Committee.

### IUE and acute brian slice imaging

*In utero* electroporation (IUE) was used to deliver the DNA encoding NIR-GECO2 and CoChR or NIR-GECO2G to the mouse brain. Briefly, embryonic day (E) 15.5 timed-pregnant female Swiss Webster (Taconic) mice were deeply anesthetized with 2% isoflurane. Uterine horns were exposed and periodically rinsed with warm sterile PBS. A mixture of plasmids pAAV-CAG-NIR-GECO2-WPRE and pCAG-CoChR-mTagBFP2-Kv2.2motif-WPRE or plasmid pAAV-CAG-NIR-GECO2G (at total DNA concentration ~1-2 μg/μL) were injected into the lateral ventricle of one cerebral hemisphere of an embryo. Five voltage pulses (50 V, 50 ms duration, 1 Hz) were delivered using round plate electrodes (ECM™ 830 electroporator, Harvard Apparatus). Injected embryos were placed back into the dam, and allowed to mature to delivery.

Acute brain slices were obtained from Swiss Webster (Taconic) mice at postnatal day (P) P11 to P22, using standard techniques. Mice were used without regard for sex. Mice were anesthetized by isoflurane inhalation, decapitated and cerebral hemispheres were quickly removed and placed in cold choline-based cutting solution consisting of (in mM): 110 choline chloride, 25 NaHCO_3_, 2.5 KCl, 7 MgCl_2_, 0.5 CaCl_2_, 1.25 NaH_2_PO_4_, 25 glucose, 11.6 ascorbic acid, and 3.1 pyruvic acid (339-341 mOsm/kg; pH 7.75 adjusted with NaOH) for 2 min, blocked and transferred into a slicing chamber containing ice-cold choline-based cutting solution. Coronal slices (300 μm thick) were cut with a Compresstome VF-300 slicing machine, transferred to a holding chamber with artificial cerebrospinal fluid (ACSF) containing (in mM) 125 NaCl, 2.5 KCl, 25 NaHCO_3_, 2 CaCl_2_, 1 MgCl_2_, 1.25 NaH_2_PO_4_ and 11 glucose (300-310 mOsm/kg; pH 7.35 adjusted with NaOH), and recovered for 10 min at 34 °C, followed by another 30 min at room temperature. Slices were subsequently maintained at room temperature until use. Both cutting solution and ACSF were constantly bubbled with 95% O_2_ and 5% CO_2_.

Individual slices were transferred to a recording chamber mounted on an inverted microscope (Eclipse Ti-E, Nikon) and continuously superfused (2–3 mL/min) with ACSF at room temperature. Cells were visualized through a 10× (0.45 NA) or 20× (0.75 NA) air objective with epifluorescence to identify positive cells. The fluorescence of NIR-GECO2 or NIR-GECO2G was excited by a SPECTRA X light engine (Lumencor) with 631/28 nm excitation and was collected through a 664LP emission filter, and imaged onto an Orca-Flash4.0 V2 sCMOS camera (Hamamatsu). Optical stimulation of slices expressing NIR-GECO2 and CoChR was performed using a 470 nm LED (ThorLabs, M470L3) at 0.157 mW/mm^2^. 4-AP stimulation was done by adding 4-AP solution to the imaging chamber at a final concentration of 1 mM.

### Imaging of NIR-GECO2 in *C. elegans*

Worms were cultured and maintained following standard protocols^25^. The genes of NIR-GECO2, NIR-GECO2G, heme-oxygenase 1, CoChR and jGCaMP7s for expression in *C. elegans* were codon-optimized using SnapGene codon-optimization tool and synthesized by GenScript. Transgenic worms expressing NIR-GECO2(G) and jGCaMP7s pan-neuronally or NIR-GECO2 in AVA and CoChR-GFP in ASH were generated by injecting the plasmids tag-168::NLS-NIR-GECO2(G)-T2A-HO1 and tag-168::NLS-jGCaMP7s or plasmids flp-18::NIR-GECO2-T2A-HO1, sra-6::CoChR-SL2-GFP, and elt-2::NLS-GFP into N2 background worms, respectively, picking those with the strongest expression of green fluorescence (in neurons for the pan-neuronal strain and in the gut for optogenetic strain). NLS sequence used in this experiment was PKKKRKV.

Hermaphrodite transgenic worms were picked at L4 stage of development and put onto NGM plates with freshly seeded OP50 lawns 12-24 h before experiments, with or without 100 μM ATR for optogenetic experiments. Worms were mounted on 2% agarose pads on microscope slides, immobilized with 5 mM tetramisole, covered by a coverslip, and imaged using a Nikon Eclipse Ti inverted microscope equipped with a confocal spinning disk (CSU-W1), a 40×, 1.15 NA water-immersion objective and a 5.5 Zyla camera (Andor), controlled by NIS-Elements AR software. Fluorescence of NIR-GECO2 was imaged with 640-nm excitation provided by 41.9 mW laser and 685/40 nm emission filter; jGCaMP7s/GFP fluorescence was imaged with a 488-nm excitation provided by 59.9 mW laser and a 525/50 nm emission filter. Optogenetic stimulation was performed with 488 nm illumination at 20 mW/mm^2^. For 200 mM NaCl stimulation, worms were imaged using the same optical setup as above, using a microfluidic device that was described previously^20^.

### Imaging of NIR-GECO2G in *Xenopus laevis* tadpoles

NIR-GECO2G and GCaMP6s were cloned into the pCS2+ vector and the plasmid was linearized with *NotI*. Capped mRNA of NIR-GECO2G and GCaMP6s was transcribed with the SP6 mMessageMachine Kit (Ambion, Thermo Fisher). The RNA (500 pg of each sample) was injected in one blastomere at the 2-cell stage resulting in animals expressing NIR-GECO2G and GCaMP6s protein in one lateral half of the animal. The animals were kept at 20°C until stage 47. Immediately before imaging the tadpole was paralyzed with pancuronium bromide (1.5 mg/mL in 0.1× MSBH) and embedded in 1% low-melt agarose.

Light-sheet imaging was performed on a Zeiss Z1 located in the McGill Advanced BioImaging Facility. The instrument was equipped with a sCMOS camera (1920×1920 pixel PCO.Edge), excitation lasers with wavelengths of 488 nm (75 mW max. output) and 640 nm (50 mW max. output). For acquisition, we excited the sample with 10% intensity of each wavelength through 5× LSFM excitation objectives (NA = 0.1) resulting in a 4.53 μm thick light-sheet. We utilized a water immersion objective for detection (Zeiss 10× PLAN APOCHROMAT, NA = 0.5, UV-VIS-IR, ND = 1.336) allowing the sample to be immersed in 0.1× MBSH. The fluorescence was directed via a dichroic mirror LBF 405/488/640 nm and filtered depending on the probe with 505-545 nm bandpass or 660 nm longpass emission filters.

The imaging data (**Fig. 4**) was acquired with 50 ms integration time and 2.39 seconds cycle time through the volume. The instrument has a short deadtime to home the axial position leading to a scanning frequency of 3.0 s for the entire volume. The raw data was corrected for drift and rapid movement with the ImageJ plugin TurboReg^26^. The image in **Fig. 4a** is a volume projection of a 45 μm thick volume capturing the spontaneous Ca^2+^ responses in the olfactory bulb. The cells were manually selected if they showed a spontaneous Ca^2+^ response as detected with both NIR-GECO2G and GCAMP6s (**Fig. 4b**). The fluorescence response was measured as mean fluorescence intensity per cell and normalized by I_norm_ = (I_m_-I_min_)/(I_max_-I_min_). I_m_ indicates the measured mean value per area and I_max_ and I_min_ the maximal and minimal value measured for the specific ROI. The normalized response of NIR-GECO2G and GCAMP6s of a cell is plotted over time in the graph in **Fig. 4c**.

## Data and image analysis

All images in the manuscript were processed and analyzed using either ImageJ (NIH) or NIS-Elements Advanced Research software (Nikon). Traces and graphs were generated using GraphPad prism 8, Origin (OriginLab) and Matlab. Data are presented as mean ± s.d. or mean ± s.e.m, as indicated.

## Data availability

Plasmids used in this study and the data that support the findings of this study are available from the corresponding author on reasonable request.

## Acknowledgments

We thank the University of Alberta Molecular Biology Services Unit for technical support. We thank Ahmed S. Abdelfattah from Janelia Research Campus for testing NIR-GECO variants and Panagiotis Symvoulidis from MIT for help with data analysis. We also thank Xian Xiao and Hongyun Tang from Westlake University for help with *C.elegans* imaging. K.D.P. acknowledges the Foundation of Westlake University. S.A. thanks Conrad F. Harrington Fellowship from the faculty of medicine at the University of McGill. Work in the lab of R.E.C. was supported by grants from CIHR (MOP-123514 and FS-154310), NSERC (RGPIN 288338-2010 and 2018-04364), Brain Canada, and NIH (U01 NS094246 and UO1 NS090565). E.S.B. acknowledges Lisa Yang, John Doerr, U.S.-Israel Binational Science Foundation (2014509), NIH (2R01-DA029639, 1R01-MH12297101, and 1RF1-NS113287), and NSF 1848029 and 1734870. Work in the lab of E.S.R. was supported by grants from CIHR (FDN-143238) and Brain Canada. Work in the lab of P.W.W. was supported by a grant from NSERC (RGPIN-2017-05005).

## Author contributions

Y.Q. developed the new NIR-GECO variants and performed characterizations *in vitro* and in dissociated neurons. D.O.C, O.T.C., Y.Q. and K.D.P. performed experiments on *C.elegans.* S.A. and A.S. performed experiments on *Xenopus laevis* tadpoles. Y.Q., K.D.P and M.H.M characterized NIR-GECO2 and NIR-GECO2G in acute brain slices. W.C.S and Y.F.C. performed imaging in HeLa cells and iPSC-CMs. A.A. measured stopped-flow kinetics. Y.Q. designed and characterized miRFP-based GECI. Y.F.C, P.W.W., E.R., E.S.B., and R.E.C. supervised research. Y.Q. and R.E.C. wrote the manuscript.

## Supplementary Materials

**Supplementary Table. 1.** Spectral, photochemical and biochemical properties of NIR-GECO2G, NIR-GECO2 and NIR-GECO1.

| Indicator | [Ca<sup>2+</sup>]<br>(mM) | EC ($\times 1000$<br>mM<sup>-1</sup>*cm<sup>-1</sup>) | QY (%) | pK<sub>a</sub> | K<sub>d</sub><br>(nM) | Hill<br>Coeff.<br>(n) | k<sub>off</sub><br>(s<sup>-1</sup>) |
| --- | --- | --- | --- | --- | --- | --- | --- |
| NIR-GECO2G | 0 | 74 | 6.1 | 5.34 | 480 | 0.78 | 3.678 |
|  | 5 | 21 | 2.1 | 4.84 |  |  |  |
| NIR-GECO2 | 0 | 67 | 5.9 | 5.26 | 331 | 0.94 | 2.995 |
|  | 5 | 18 | 1.4 | 4.85 |  |  |  |
| NIR-GECO1 | 0 | 69 | 6.0 | 5.14 | 885 | 0.99 | 2.354 |
|  | 5 | 20 | 2.0 | 4.84 |  |  |  |

Abbreviations: EC, extinction coefficient; QY, quantum yield; pK<sub>a</sub>, pH corresponding to 50% of the maximal fluorescence brightness measured at optimal pH; K<sub>d</sub>, K<sub>d</sub> for Ca<sup>2+</sup>; k<sub>off</sub>, Ca<sup>2+</sup>-dissociation kinetics measured by stopped-flow spectrometer.

**Supplementary Figure 1.**
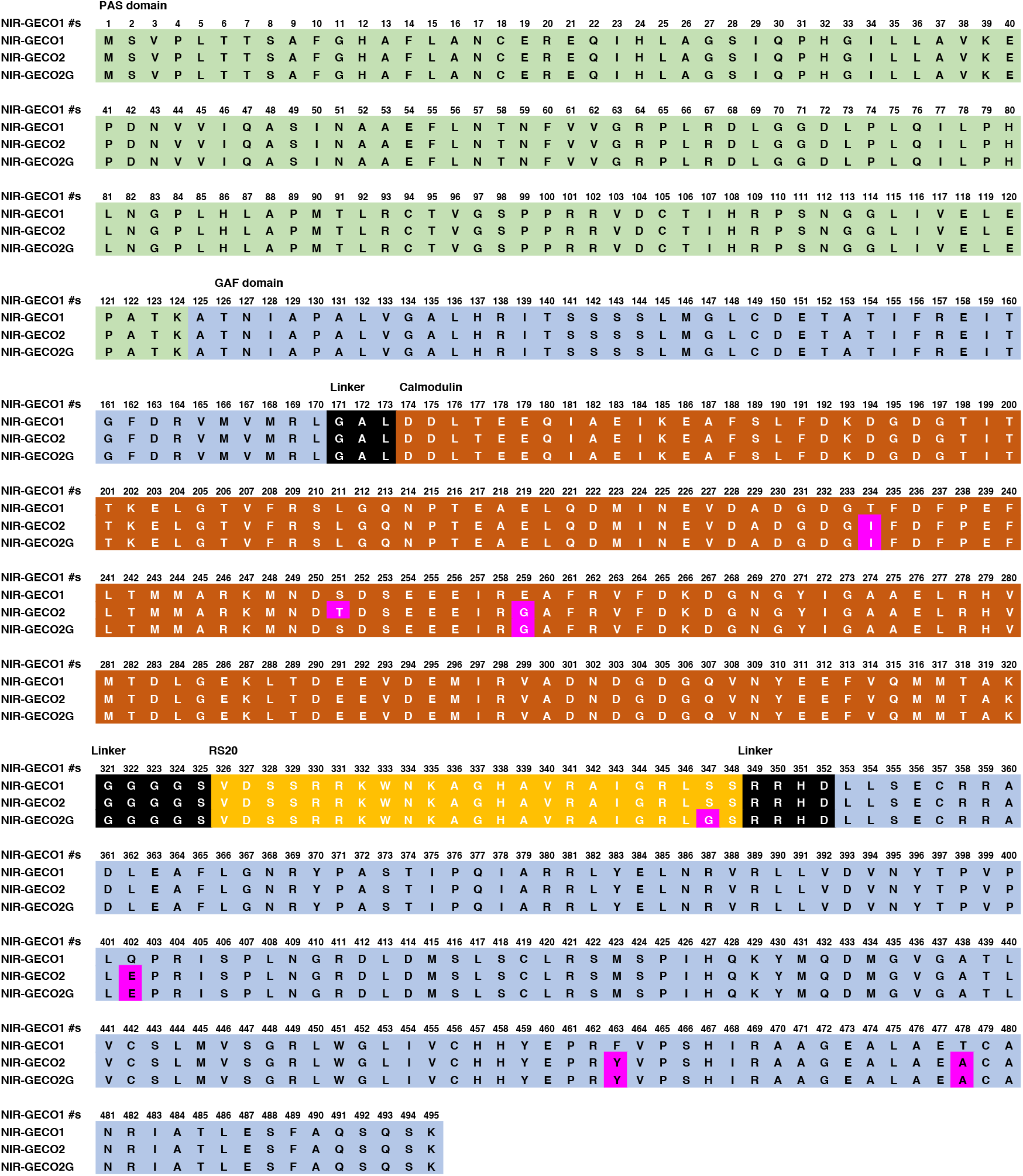
Sequence alignment of NIR-GECO2, NIR-GECO2G and NIR-GECO1. Single-amino-acid changes relative to NIR-GECO1 are highlighted with a magenta background. PAS domain, GAF domain, linkers, calmodulin, and RS20 are shown as light green, light blue, black, brown and yellow, respectively.

**Supplementary Figure 2.**
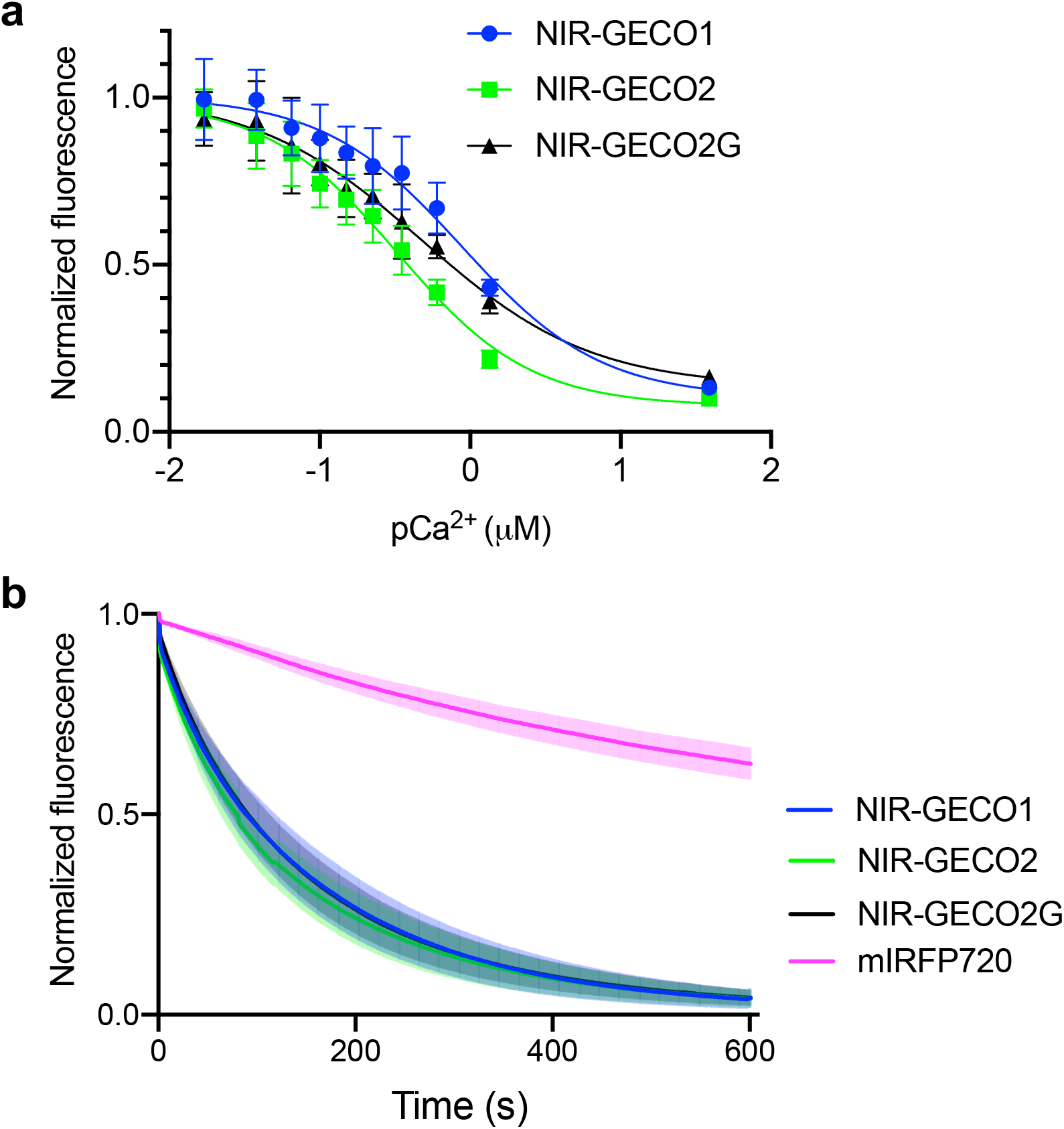
Additional *in vitro* characterization of NIR-GECO variants. (**a**) Ca^2+^ titration curves of NIR-GECO1, NIR-GECO2 and NIR-GECO2G (center values are the mean and error bars are s.d.; n = 3). **(b)** Photobleaching curves of NIR-GECO variants and mIRFP720 (n = 11, 11, 16 and 9 neurons for NIR-GECO1, NIR-GECO2, NIR-GECO2G and miRFP720, respectively. Mean value (solid line) and s.d. (shaded areas) are shown. Cells were continuously illuminated with 631/28 nm at 80 mW/mm^2^ during the experiment. Image was taken every 5 s.

**Supplementary Figure 3.**
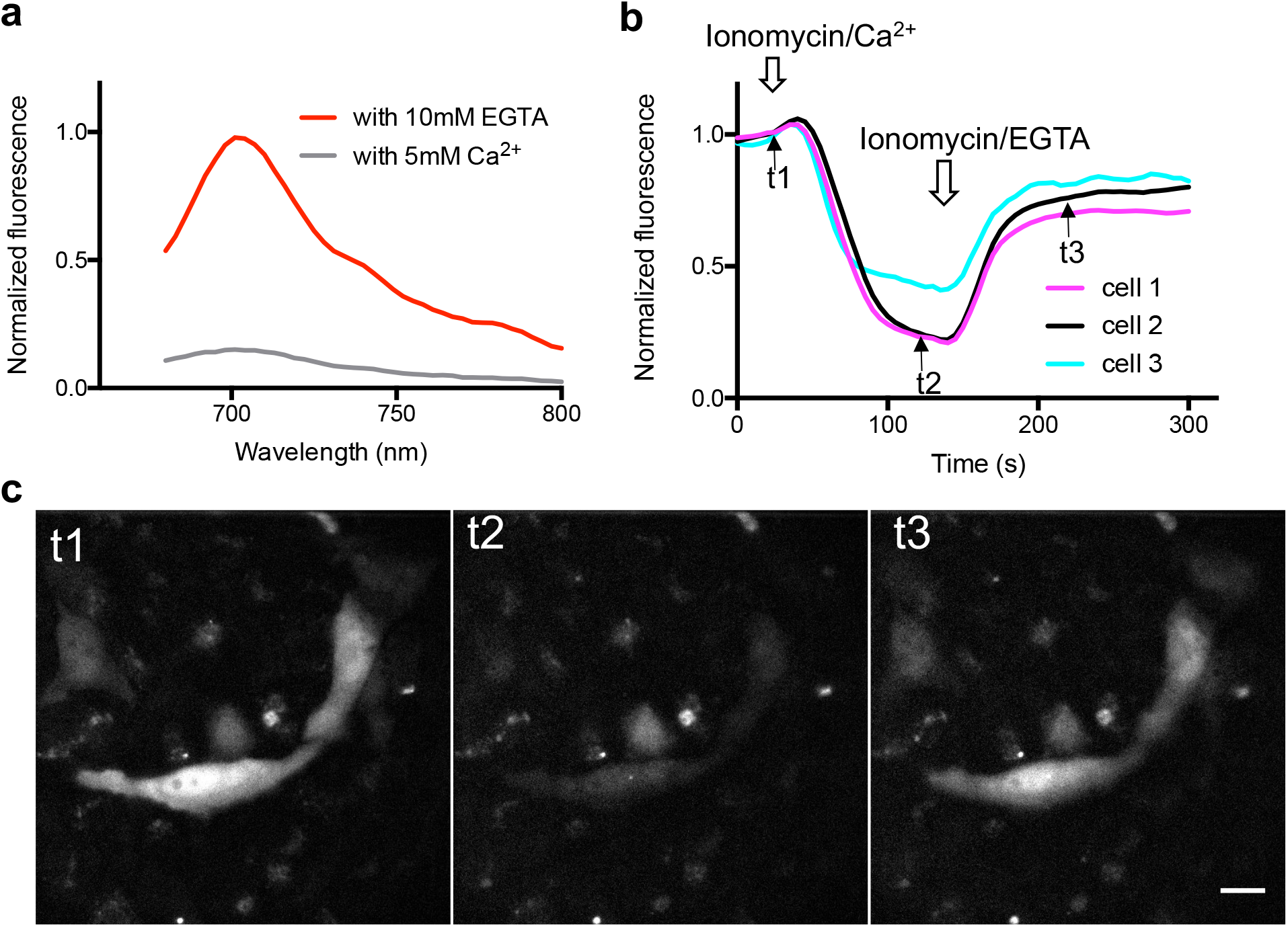
Ca^2+^ indicator prototype based on miRFP. The mIFP domain of NIR-GECO1 was replaced with miRFP using the same insertion point and linker sequences (that is, CaM-RS20 was used to replace residues 170-177 of miFP or residues 172-179 of miRFP, **Supplementary Fig. 4**). (**a**) Fluorescence emission spectra of the prototype miRFP-based Ca^2+^ indicator in the presence (5 mM Ca^2+^), and absence (10 mM EGTA), of Ca^2+^. (**b**) Intensity vs. time traces for transfected HeLa cells. Cells were treated with ionomycin/Ca^2+^ to increase cellular Ca^2+^ concentrations and ionomycin/EGTA to deplete cellular Ca^2+^. (**c**) Fluorescent images of miRFP-based Ca^2+^ indicator prototype at time points t1 to t3 (as denoted in **b**). Scale bar, 20 μm; λ_ex_=650/60 nm and λ_em_=720/60 nm. Acquisition rate 0.2 Hz. In an effort to develop an improved NIR GECI, we found that the mIFP (engineered from PAS and GAF domain of *Bradyrhizobium bacteriophytochrome*) domain of NIR-GECO1 could be replaced with miRFP^12^. miRFP is another monomeric BV-FP that was derived from *Rps. palustris bacteriophytochrome* and shares 57% amino acid homology with mIFP (**Supplementary Fig. 1, 2**). In principle, the miRFP version on NIR-GECO1 could have served as a template for making improved NIR fluorescent Ca^2+^ indicators due to its higher brightness in mammalian cells relative to mIFP^12^. However, we decided to start our further directed evolution efforts from NIR-GECO1 for two reasons. The first reason is that NIR-GECO1 was already optimized and worked well in brain slices, and so starting from it might save time and lower risk. The second reason is that over-expression of the miRFP-based Ca^2+^ indicator appeared to be toxic to bacteria and it was challenging for us to incorporate the construct into the bacteria-HeLa screening system that we used for engineering NIR-GECO1 ^8^.

**Supplementary Figure 4.**
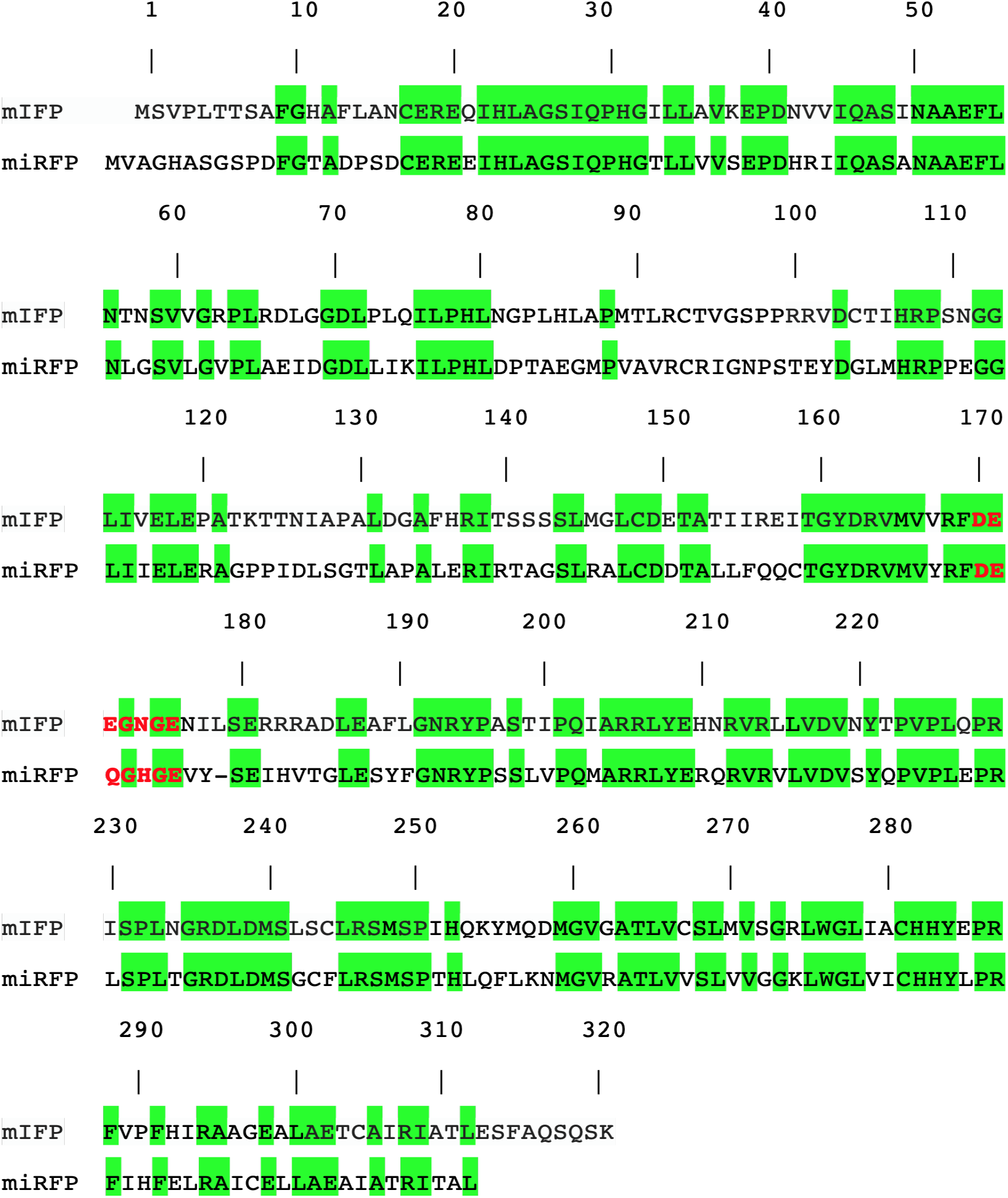
Alignment of amino acid sequences of mIFP and miRFP. Alignment numbering is based on mIFP. The structurally analogous residues between mIFP and mIRFP are highlighted in green. Residues that were replaced by CaM-RS20 to make NIR-GECO1, and the prototype mIRFP-based Ca^2+^ indicator, respectively, are in bold and red.

**Supplementary Figure 5.**
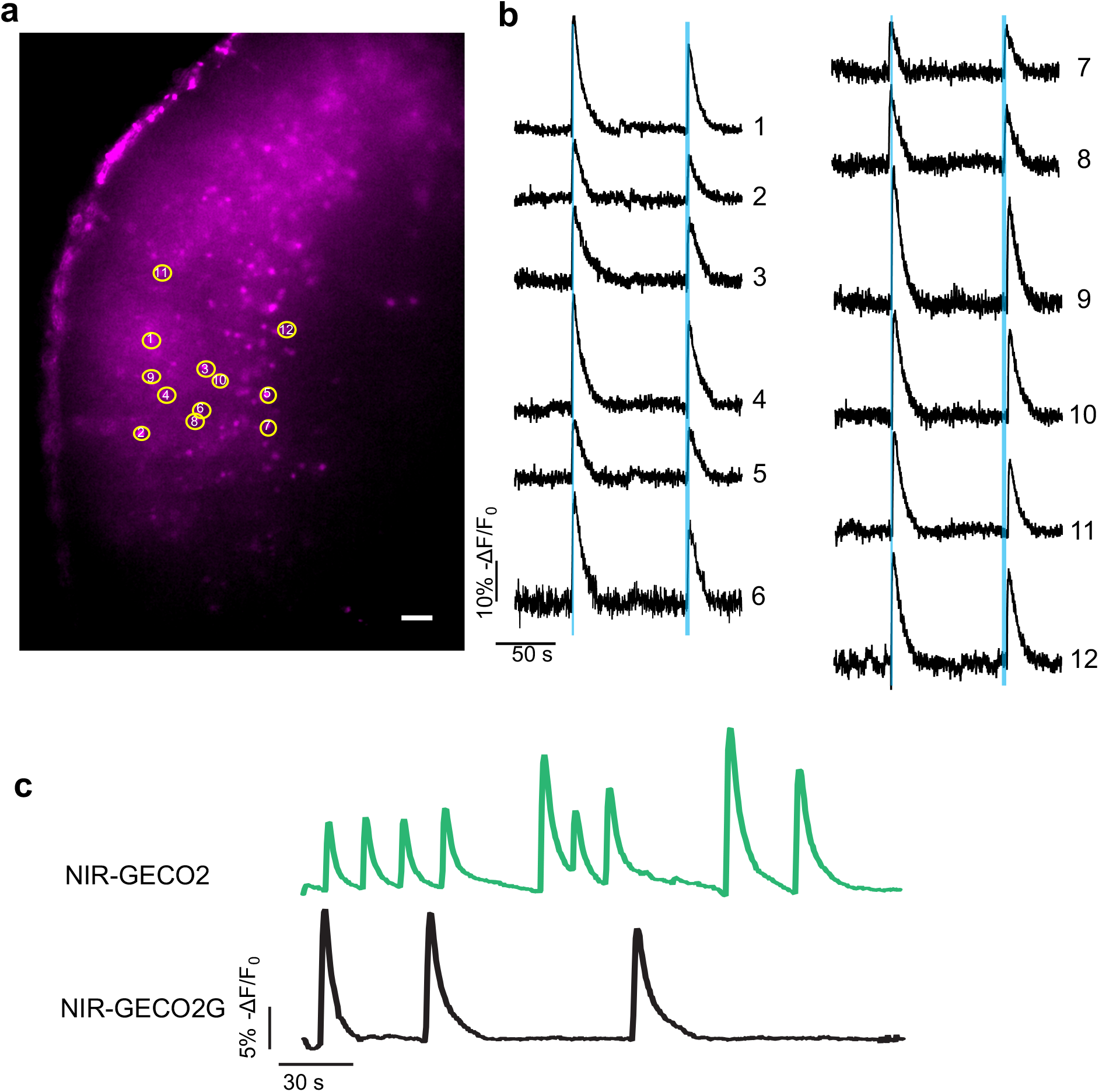
Imaging of NIR-GECO2 and NIR-GECO2G in acute brain slices. (**a**) Wide-field image of a mouse brain slice with coexpression of NIR-GECO2 and CoChR, fluorescence of NIR-GECO2 is shown (λ_ex_ = 631/28 nm and λ_em_ = 664LP). Scale bar, 50 μm (**b**) Fluorescence of NIR-GECO2 (acquisition rate 100 Hz) in response to 200 ms blue-light stimulation (470/20 nm, 0.157 mW/mm^2^, indicated by blue bar). The numbers of the traces correspond to the neurons labeled in **a**. (**c**) Single-trial wide-field imaging of 4-aminopyridine (1 mM final concentration) evoked neuronal activity from the two representative neurons in brain slices expressing NIR-GECO2 and NIR-GECO2G, respectively (λ_ex_ = 631/28 nm and λ_em_ = 664LP; acquisition rate 10 Hz).

**Supplementary Figure 6.**
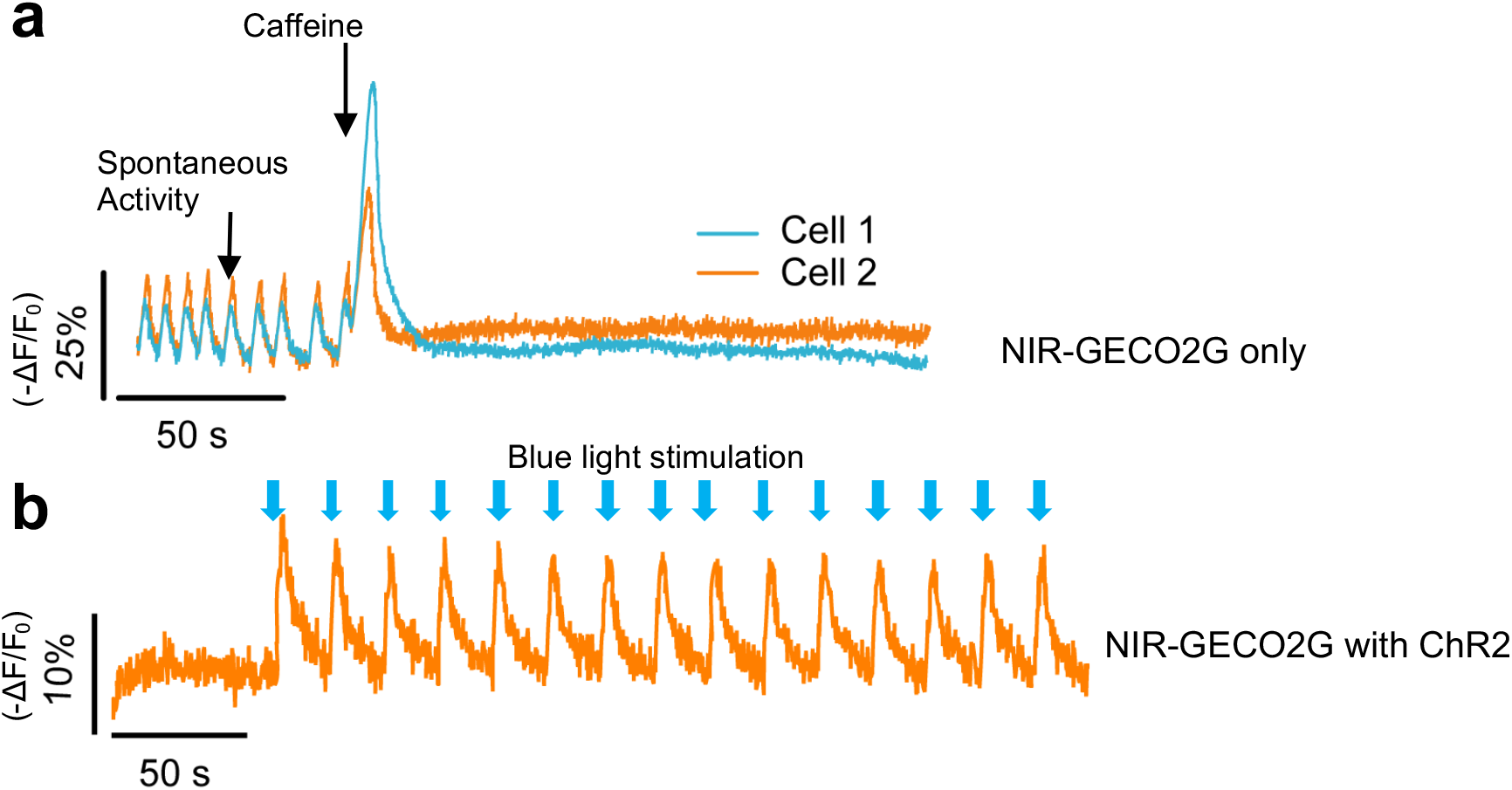
Ca^2+^ imaging in iPSC-CMs using NIR-GECO2G. (**a**) Representative fluorescence recording of spontaneous and caffeine-evoked Ca^2+^ oscillations using NIR-GECO2G in iPSC-CMs. (**b**) Representative single-trial fluorescence recording of blue-light (470 nm at a power of 1.9 mW/mm^2^) stimulated Ca^2+^ oscillations using NIR-GECO2G in channelrhodopsin-2 (ChR2) expressed iPSC-CMs.

**Supplementary Figure 7.**
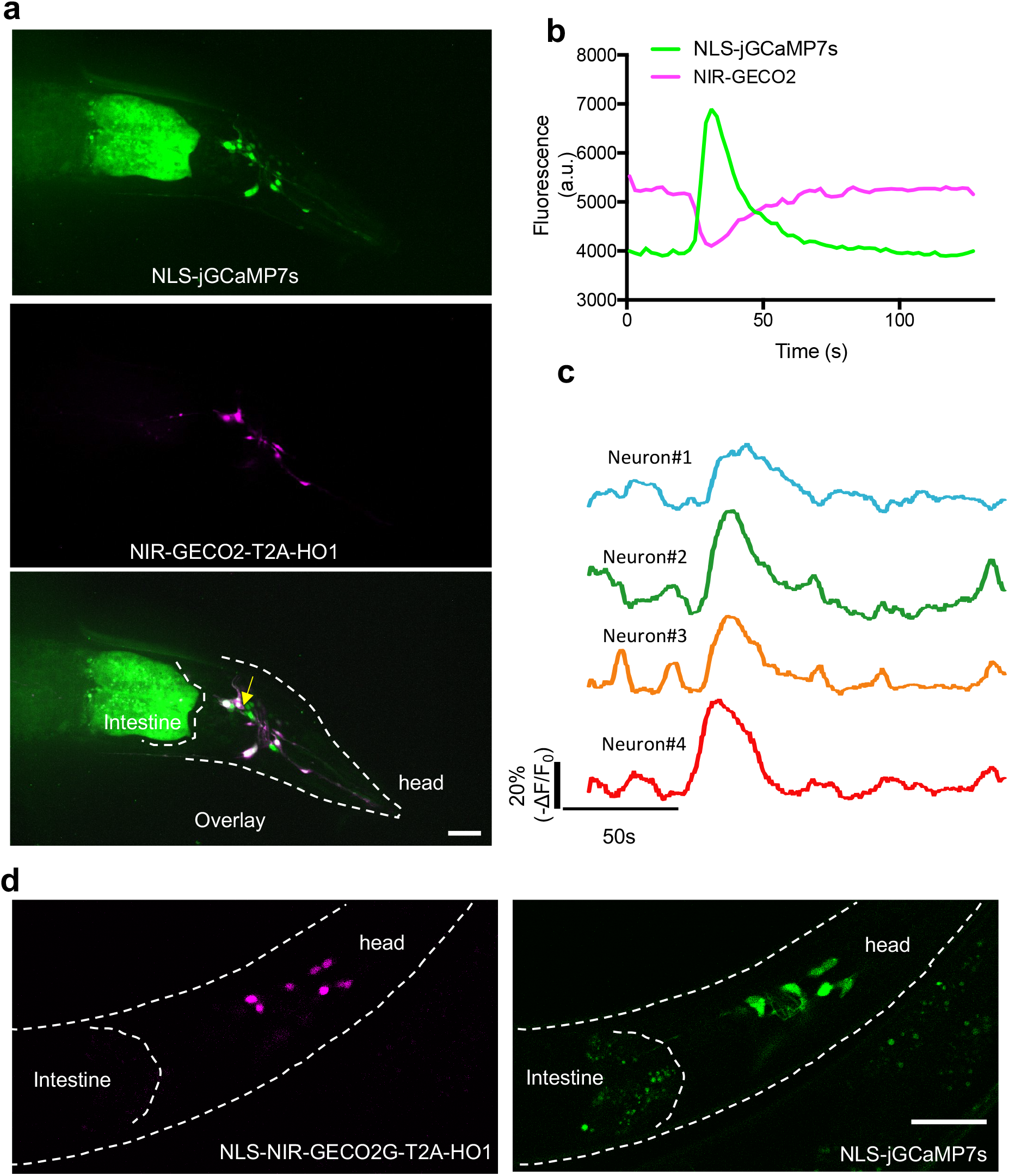
Imaging of *C. elegans* using NIR-GECO2, NIR-GECO2G and jGCaMP7s (**a**) Representative confocal images of worms co-expressing NIR-GECO2-T2A-HO1 and NLS-jGCaMP7s (representative of more than 3 worms). Top, fluorescent image of neurons expressing NLS-jGCaMP7s (λ_ex_ = 488 nm laser light, λ_em_ = 527/50 nm). Middle, fluorescent image of neurons expressing NIR-GECO2-T2A-HO1 (λ_ex_ = 640 nm laser light, λ_em_ = 685/40 nm). Bottom, Overlay image of green channel and NIR channel. Scale bar, 25 μm. (**b**) Spontaneous Ca^2+^ fluctuation of a representative worm neuron (indicated in **a** by a yellow arrow) coexpressing NIR-GECO2 and NLS-jGCaMP7s. Imaging conditions were identical to the experiments in **a**, acquisition rate 2 Hz. (**c**) Representative spontaneous Ca^2+^ oscillations of worm neurons reported by NIR-GECO2 (acquisition rate 2 Hz). Imaging conditions were identical to the experiments in **a**. (**d**) Representative confocal images of worms co-expressing NLS-NIR-GECO2G-T2A-HO1 (left) and NLS-jGCaMP7s (right). Imaging conditions were identical to the experiments in **a.** Representative of more than 3 worms. Scale bar, 25 μm.

### Supplementary Video 1 (Associated to Fig.4)

Imaging of spontaneous neuronal activity with NIR-GECO2G in the olfactory bulb of Xenopus laevis.

